# Lupus Gut Microbiota Transplants Cause Autoimmunity and Inflammation

**DOI:** 10.1101/2021.08.15.456375

**Authors:** Yiyangzi Ma, Ruru Guo, Yiduo Sun, Xin Li, Lun He, Zhao Li, Gregg J. Silverman, Guobing Chen, Feng Gao, Jiali Yuan, Qiang Wei, Mengtao Li, Liangjing Lu, Haitao Niu

## Abstract

**Background:** The etiology of systemic lupus erythematosus (SLE) is multifactorial. Recently, growing evidence suggests that the microbiota plays a role in SLE, yet whether gut microbiota participates in the development of SLE remains largely unknown. To investigate this issue, we carried out 16s rDNA sequencing analyses in a cohort of 18 female un-treated active SLE patients and 7 female healthy controls, and performed fecal microbiota transplantation from patients and healthy controls to germ-free mice.

**Results:** Compared to the healthy controls, we found no significant different microbial diversity but some significantly different species in SLE patients including *Turicibacter genus* and other 5 species. Fecal transfer from SLE patients to germ free (GF) C57BL/6 mice caused GF mice to develop a series of lupus-like phenotyptic features, which including an increased serum autoimmune antibodies, and imbalanced cytokines, altered distribution of immune cells in mucosal and peripheral immune response, and upregulated expression of genes related to SLE in recipient mice that received SLE fecal microbiota transplantation (FMT). Moreover, the metabolism of histidine was significantly altered in GF mice treated with SLE patient feces, as compared to those which received healthy fecal transplants.

**Conclusions:** Overall, our results describe a causal role of aberrant gut microbiota in contributing to the pathogenesis of SLE. The interplay of gut microbial and histidine metabolism may be one of the mechanisms intertwined with autoimmune activation in SLE.

## Introduction

In recent years the role of gut microbiota in diseases and health of hosts has drawn considerable attention. Growing evidence suggests a link between gut microbiota and various diseases including cancer [1], atherosclerosis [2], autoimmune diseases [3] and other conditions. A number of specific microbial biomarkers have been described and fecal microbiome targeted strategies are potential diagnosis and treatment of different diseases.

Systemic lupus erythematosus (SLE) is a prototypic autoimmune disease affecting multiple organs, particularly in young females. SLE is characterized by autoantibody production and dysregulation of both innate and adaptive immune systems. The combination of genetic and environmental factors is thought to be involved in the etiology of SLE, but the exact pathogenic mechanisms remain elusive [4]. Studies found that gut microbiota in SLE patients showed significantly different compared with healthy cohorts, the Firmicutes-to-Bacteroidetes ratio can be reduced [5] or not significantly different [6]; the diversity index of microbial communities are reduced in SLE [7]. The relative abundance of some species such as Lactobacillus differs from MRL/lpr [8] or NZB/W F1 mice [9]. In murine lupus models, studies have also emphasized a strong correlation between gut microbiota and SLE. Recently, it has been suggested that elimination of gut bacteria can reduce SLE pathology in some mouse models of lupus [10, 11], which indicated the gut microbiota may be potential to progression of SLE. In our previous study, we have demonstrated fecal microbiota from lupus murine model induced inflammatory response which promoted increased anti-dsDNA antibody levels in germ-free mice[12]. However, evidence is still needed to determine whether altered gut microbiota communities are a consequence or an important causal factor for the pathogenesis of SLE. Fecal transplantation from human samples into germ free mice is required to clarify the involvement of gut microbiota in pathophysiology of SLE. These issues are major goals of present studies.

To address the questions above, we performed 16S rRNA sequencing of 18 female SLE patients and 7 healthy control; transplanted fecal microbiota into germ-free C57/B6J non-autoimmune mice and evaluated for autoimmune antibodies, serum cytokines and distribution of immune cells in mucosal and peripheral immune response; took transcriptome sequencing for gene expression functional analysis and metabolism phenotype analyses of the FMT recipients.

## Results

### 1. Characterization of microbiota in SLE patients and Healthy donors

In total, 7 healthy controls and 18 un-treated SLE patients were included in this cross-sectional study, the clinical records were listed in Supplementary table S1. To investigate the community compositions and differences between SLE patient (SLE) group and healthy control (HC) group, we compared operational taxonomic units (OTUs) numbers, alpha diversity, beta diversity and LDA Effect Size (LEfSe) between groups. Fig S1 A-B showed the microbial composition of SLE patients and healthy controls. Fig S1 C-D showed the PCA analysis and Anosim results of HC group and SLE group. These two beta diversity indices indicate that the composition of the communities in the two groups was different although it was not significantly different. Fig 1 displayed the significantly different microbial species between HC group and SLE group. T-test results showed one genus (*Turicibacter*) and 5 species (*Clostridium*_*papyrosolvens, Lactobacillus_reuteri, Lactobacillus_intestinalis, Lachnospiraceae_bacteriumA2 and Lachnospiraceae_bacteriumM18-1*) were significantly different between HC group and SLE group (Fig 1A and Fig 1B). LEfSe results showed the biomarker species with a LDA score over 4 (Fig 1C), there were 5 microbial communities (*Clostridia, Clostridiales, Lachnospiraceae, unidentified_Lachnospiraceae, Subdoligranulum*) significantly enriched in the HC group from phylum to genus level, which indicated that these biomarker microbial species were related to maintain the healthy state of healthy control group. Fig 1D was the biomarker communities in species level, the results showed that non-dominant species communities may contribute much to SLE.

**Figure 1.**
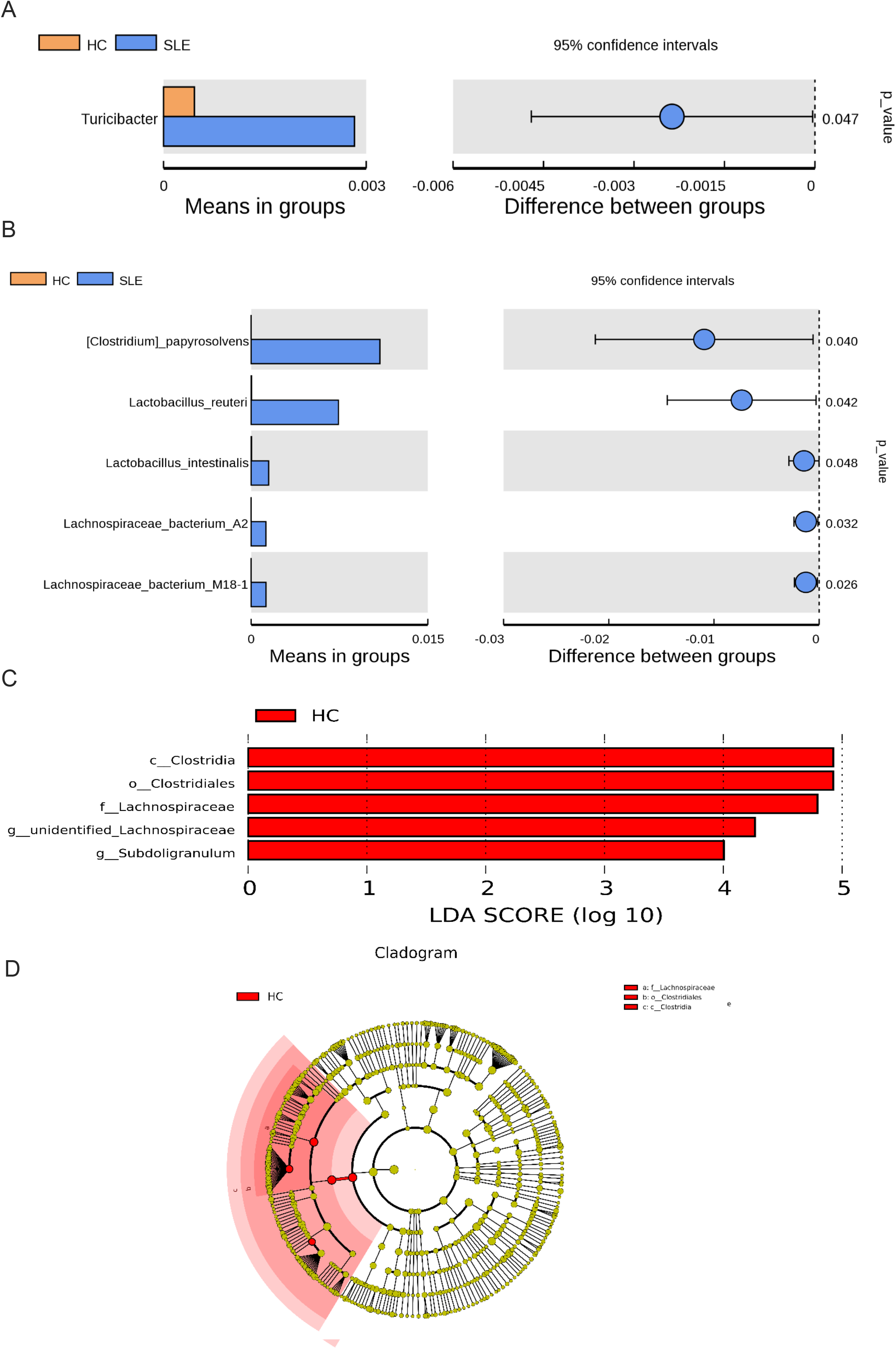
Analysis of significantly different microbial communities in SLE patients and healthy control. Feces of each group were collected for analysis of bacterial 16S RNA gene amplicon libraries and determination of the different species. Comparison of different species between groups in genus(B) and species (B) by t-test. LEfSe analysis of the different microbial communities between groups. LDA scores for different bacterial taxa of the two groups from phylum to genus(C). LDA scores for different bacterial taxa of the two groups in species (D). (p, phylum; c, class; o, order; f, family; g, genus)

### 2. Microbiota of human SLE inhabits in intestinal of germ-free mice with similar profile

We transplanted the feces obtained from 4 SLE patients and 4 healthy people separately into different germ free mice. To clarify the colonization of human fecal microorganisms in recipient mice, at the 3^rd^ week after transplantation we collected individual fecal samples and analyzed these communities by 16s rRNA amplicon sequencing. Fig S2A-B displayed the microbiota composition of the recipient mice at the phylum and genus level. Fig 2A and Table 1 showed the Source tracker analysis results of the recipient mice. The results indicated that microbiota from healthy control contribute 76.14% (±7.23) within GF+HC group; microbiota from SLE patients contribute 83.00% (±3.28) within the GF+SLE group. Fig S2 (C, D) showed PCA and Anosim index between recipients groups. Fig 2B illustrates the species that were significantly different between these groups.

**Figure 2.**
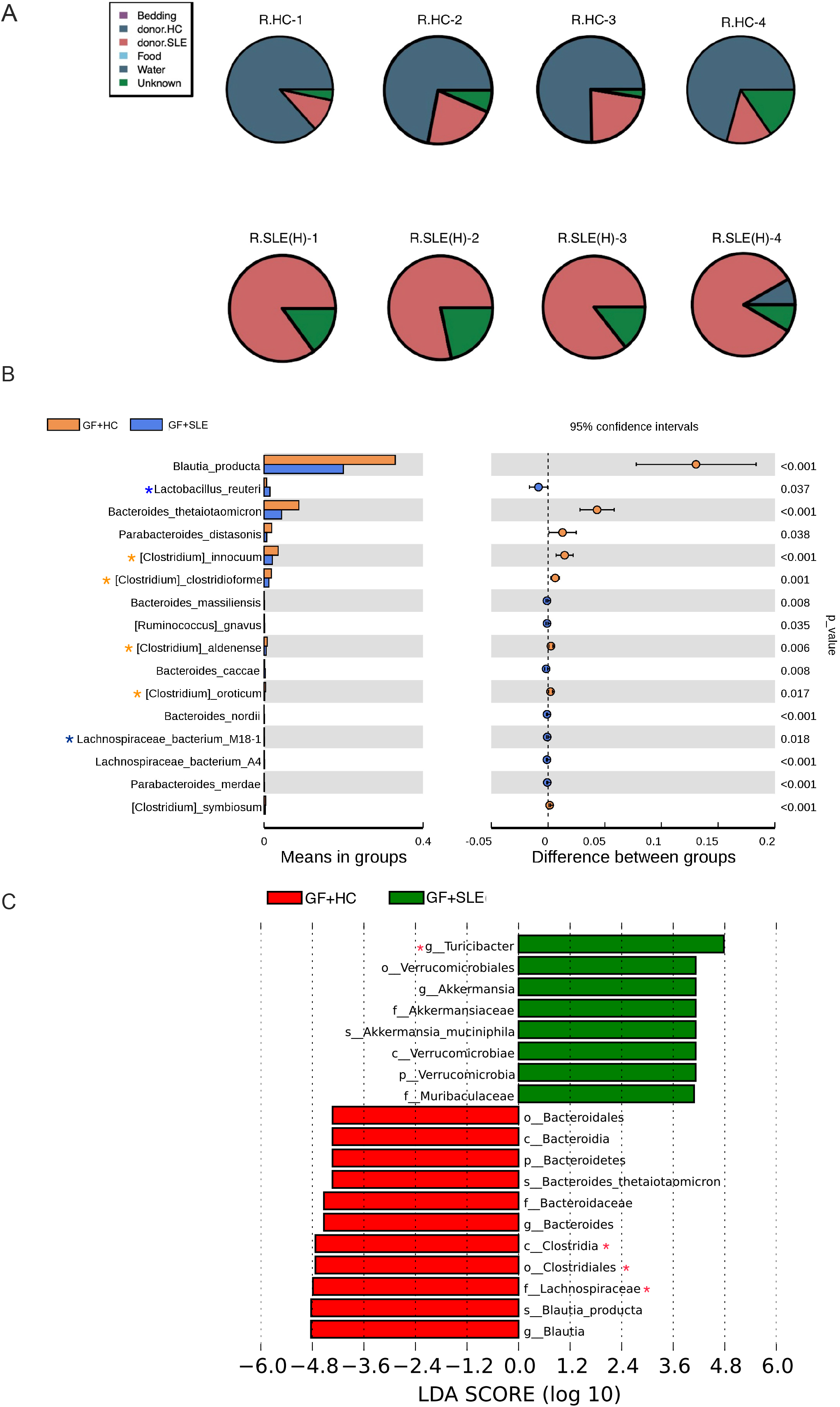
FMT from SLE patients and healthy control into GF mice. (A) SourceTracker results of recipient mice (4 mice per group). The first row shows the result of each mouse in the GF+HC group and the second row shows the result of each mouse in the GF + SLE group. (B) Significantly different genus in GF+SLE group and GF+HC group (4 mice per group). Yellow symbol* in the left column showed the genus of GF+SLE in accordance with SLE patients; Blue symbol* in the left column showed the genus of GF+HC in accordance with healthy control; (C) LDA scores for different bacterial taxa of the two recipients’ groups in species (D). (p, phylum; c, class; o, order; f, family; g, genus)

**Table. 1.** SourceTracker results for mice in the GF + HC group and the GF + SLE group.

| Group | Sample | Bedding (%) | HC(%) | Food (%) | SLE(%) | Water (%) | Unknown (%) |
| --- | --- | --- | --- | --- | --- | --- | --- |
| GF+HC | R.HC.1 | 0 | 86.59 | 0 | 10.11 | 0 | 3.29 |
|  | R.HC.2 | 0 | 71.97 | 0 | 21.43 | 0 | 6.60 |
|  | R.HC.3 | 0 | 75.29 | 0 | 22.20 | 0 | 2.51 |
|  | R.HC.4 | 0 | 70.71 | 0 | 13.76 | 0 | 15.47 |
| GF+SLE | R.SLE(H).1 | 0 | 0 | 0 | 85.01 | 0 | 14.99 |
|  | R.SLE(H).2 | 0 | 0 | 0 | 78.31 | 0 | 21.69 |
|  | R.SLE(H).3 | 0 | 0 | 0 | 85.51 | 0 | 14.42 |
|  | R.SLE(H).4 | 0 | 8.30 | 0 | 83.16 | 0 | 8.54 |

Specially, *Clostridium innocuum, Clostridium clostridioforme, Clostridium aldenense and Clostridium oroticum,* were significantly enriched in GF+HC group; *Lactobacillus_reuteri* was significantly enriched in GF+SLE group (Fig 2B). What is inreststing about the LEfSE results showed in Fig 2C is the *Turicibacter Clostridia, Clostridiales, Lachnospiraceae.* These results were as same as tendency of human donors. These finding document that following fecal microbiota transplantation the fecal microbiota of human beings can colonize the intestines of germ-free mice, and the specific composition of the community within the recipient mice is largely dependent on the human donors.

### 3. Microbiota of human SLE causes increased autoantibody levels and abnormal inflammatory responses in germ-free mice

The serum of recipients mice after FMT were collected and the titers of IgG anti-dsDNA antibody were tested by ELISA each week. From Fig 3A we can see that the level of anti-dsDNA antibody increased and there was a significantly higher level of anti-dsDNA antibodies in GF+SLE group at 2 weeks and 3 weeks after FMT. The recipients were sacrificed at 3 weeks after FMT, when we detected anti-dsDNA antibody (Fig 3B), anti-ANA antibody (Fig 3C) and total IgG antibodies levels (Fig 3D) in their sera. The content of anti-dsDNA antibodies and anti-ANA antibodies in GF+SLE group were both significantly higher than GF+HC group; there was no significant differences in the content of total IgG in the GF+SLE group as compared with the GF+HC group. These results indicated that GF mice that received SLE patients fecal microbiota developed higher autoimmune antibody levels, which were considered to a close pathologic feature of SLE. We detected total IgA in feces and some inflammatory cytokine in serum of recipient mice. Fecal IgA levels in healthy donors were significantly higher than that in SLE donors (Fig 3D). The IgA levels in recipient mice with different fecal microbiota transplantation were both significantly increase while the content in feces between recipients groups were similar (Fig 3F).

**Figure 3.**
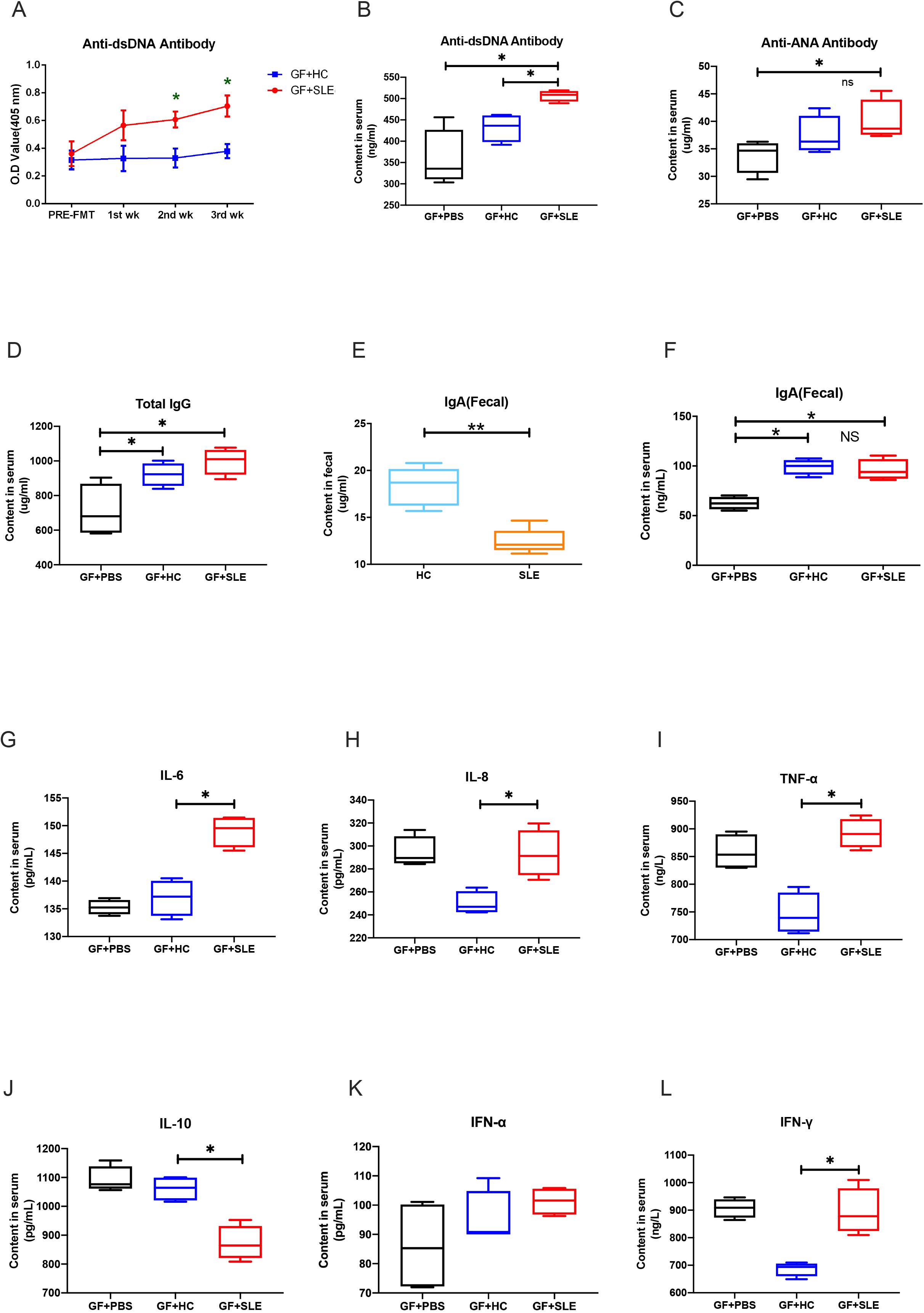
Analysis of immune response in recipient intestinal and serum. (A) Anti-dsDNA antibody titers of mice before FMT to 3 weeks after FMT in the GF+SLE group and the GF+HC group (4 mice per group). (B) Content of anti-dsDNA antibody in serum of GF+PBS group, GF+HC group and GF+SLE group. (C) Content of anti-ANA antibody in serum of GF+PBS group, GF+HC group and GF+SLE group. (D) Content of total IgG in 3 recipient mice groups. (E) Content of IgA in feces from SLE patients and healthy control. (F) Content of IgA in feces from 3 recipient mice groups. (G-J) Content of cytokine IL- 6, IL-8, TNF-α and IL-10 in serum of 3 recipient mice groups. (K-L) Content of IFN-α and IFN-γ in serum of 3 recipient mice groups. Data represent one of three independent experiments, and error bars represent means ± SEMs. \**p* < 0.05, \*\**P* < 0.01.

Fig 3 G-L presents the murine cytokine levels in recipient mouse sera. IL-6, IL-8, TNF-α, and IFN-γ were all significantly increased in GF+SLE group as compared with GF+HC group. a higher level of inflammation response in recipients that received SLE patients fecal (Fig 3G-I, L). There was no difference between groups for IFN-α (Fig 3K). Notably, IL-10 as a regulatory cytokine, was significantly decreased in the GF+SLE mice (Fig 3J). These results suggest that GF mice which received fecal microbiota from SLE patients produced a higher pro-inflammation and a decreased immune regulatory response.

### 4. Microbiota of human SLE influences the distribution of lymphocytes in colonized germ-free mice in mucosal and peripheral immune system

Lamina propria (LP) are essential for immune responses in intestinal mucosal immunity. We examined the lymphocytes distributions in LP of mice receiving different treatments. There were no detectable B cell subsets due to the lack of microbiota stimulation in GF+PBS group (data not shown). We analyzed B cells and its subsets in fecal transplanted mice. In LP, CD19^+^ B cells and macrophages were significantly increased in GF+SLE group compared with GF+HC group (Fig 4A-B). The frequency of plasma cells were significantly increased in GF+SLE group (Fig 4C). The germinal center B cells, immature B cells were elevated in GF+SLE group (ns, Fig 4S, A-B). Due to the paucity of T cells in the GF mice, the representation of CD4^+^ T cell subsets. Recipient mice in GF+HC group and GF+SLE group had increased numbers of CD4^+^ T cells and its subsets as compared with GF+PBS group, although there were no significant difference in these two groups (Fig 4S, C).

**Figure 4.**
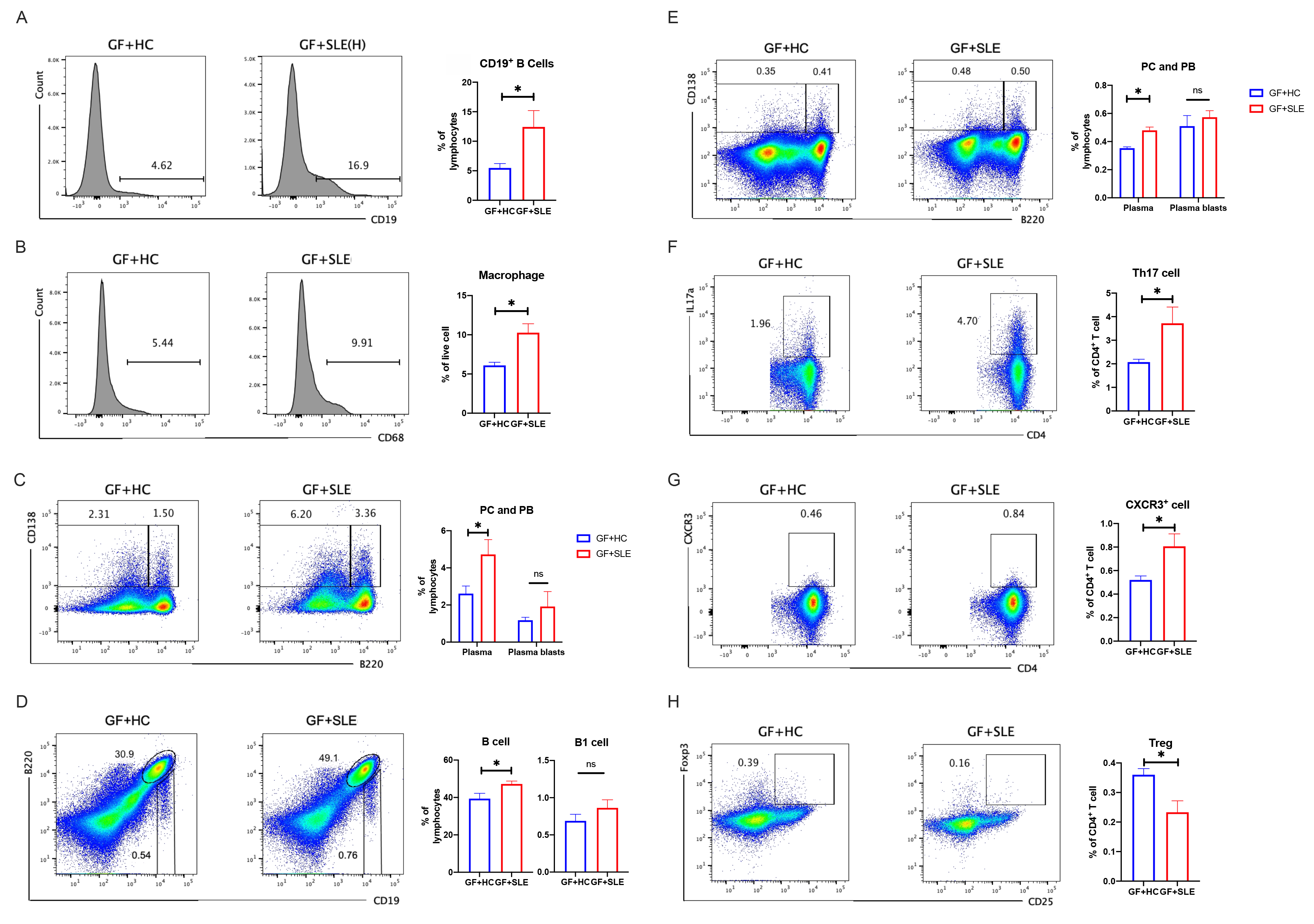
Flow cytometry analysis of the distribution of lymphocytes in the lamina propria (LP) and spleen (SP) of GF+HC mice and GF+SLE mice. (A) Plots of B cells (left) and percentages of B cells in LP (4 mice per group, right). (B) Plots of macrophage (left) and percentages of macrophage in LP (4 mice per group, right). (C) Plots of plasma cells (PCs) and plasma blast cells (PBs;left) and percentages of PCs and PBs in LP (4 mice per group; right). (D) Plots of B cells and B1 cells (left) and percentages of B cells and B1 cells in spleen (4 mice per group, right). (E) Plots of plasma cells (PCs) and plasma blast cells (PBs;left) and percentages of PCs and PBs in SP (4 mice per group; right). (F) Plots of Th17 cells (left) and percentages of Th17 cells in SP (4 mice per group, right). (G) Plots of CD4+CXCR3+ cells (left) and percentages of CD4+CXCR3+ cells in SP (4 mice per group, right). (H) Plots regulatory T cells (Tregs, left) and percentages of Tregs in SP (4 mice per group, right). Data represent one of two independent experiments. A Mann-Whitney t-test was used to compare groups, and error bars represent means ± SEMs. \**P* < 0.05, \*\**P* < 0.01, \*\*\**P* < 0.001.

We also examined the status of the peripheral immune organ after transplantation for measurement of lymphocyte distributions in the spleen. Fecal microbiota transplantation increased the frequency of B and T cells, as well as its subsets in both GF+HC and GF+SLE group. We observed an elevation of B cells and B1 cells (Fig 4D), Plasma cells and plasma blasts (Fig 4E) immature B cells (Fig 4S,D), transitional B cells (Fig 4S,E), among these results we can see that the proliferation of B cells, immature B cells and plasma cells were much more increased in GF+SLE group(p<0.05, Fig). We also detected the distribution of some T cell subsets. There were increased CD4^+^ and CD8^+^ T cells in both fecal microbiota treated groups (Fig 4s,C), specially, the frequency of Th17 cells and CD4^+^CXCR3^+^ cells were increased significantly in GF+SLE group(p<0.05, Fig4F,4G); CD25^+^Foxp3^+^ Treg cells were decreased in GF+SLE group(p<0.05, Fig 4H). Together these results indicated that SLE fecal microbiota had a greater effect of inflammation response in recipients mice compared with HC fecal microbiota.

### 5. Microbiota of human SLE impacts the differential gene expression and enrichment in colonized germ-free mice

We determined the effect of FMT from SLE patients and healthy donors on the differential expression of genes (DEGs) and their enrichment by transcriptome sequencing in recipients’ intestines and spleens. We analyzed the DEGs in recipients intestines. Comparison of GF mice that received microbiota from SLE with the mice received microbiota from healthy donors, patients indicated there were 1085 DEGs(based on a threshold change of 1.3-fold or more, and adjusted for a q-value less than 0.05), with 688 upregulated genes and 397 downregulated genes (Fig 5A); these DEGs were mainly enriched in retinol metabolism, steroid hormone biosynthesis, tryptophan metabolism, fatty acid degradation and some other metabolism pathways (Fig 5A). Additionally, DEGs enrichment results in GF+SLE versus GF+PBS mainly concentrated in metabolism as showed in Fig 5S.

**Figure 5.**
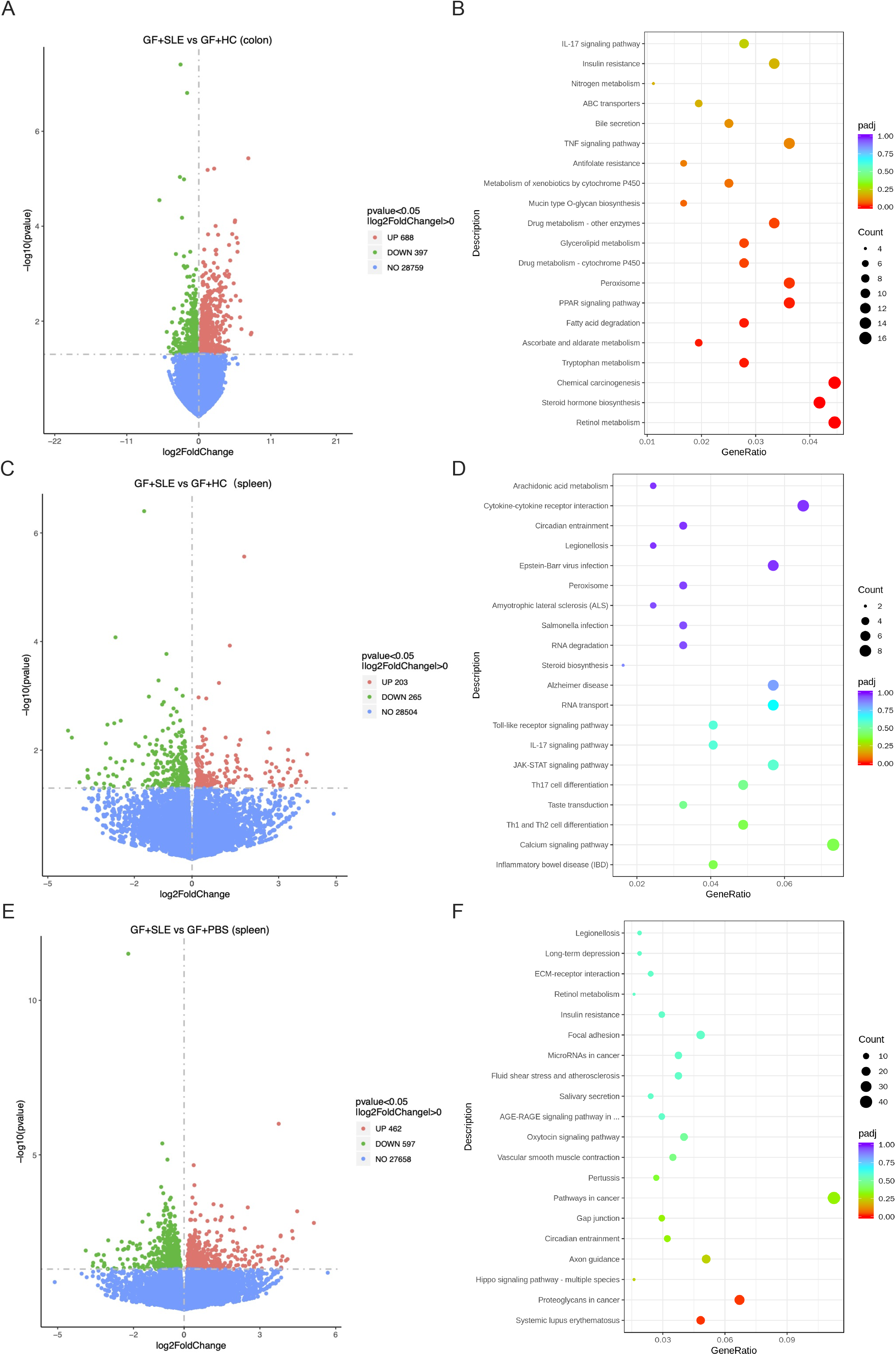
Differential expression of genes (DEGs) in the intestines and spleen of GF mice after different treatments. (A) Comparison of the GF + SLE and the GF + HC groups. The numbers of differential expressed genes in colons showed in volcano plot (left) and its enrichment in KEGG (right). (B) Comparison of the GF + SLE and the GF + HC groups. The numbers of differential expressed genes in spleen showed in volcano plot (left) and its enrichment in KEGG (right). (C) Comparison of the GF + SLE and the GF + PBS groups. The numbers of differential expressed genes in spleen showed in volcano plot (left) and its enrichment in KEGG (right).

The DEGs in recipients’ spleen were also analyzed. Comparison of GF mice that received microbiota from SLE patients indicated there were 468 DEGs, with 203 upregulated genes and 263 downregulated genes; the functions of these DEGs showed that DEGs participants in pathways including inflammatory bowel disease (IBD), Th1 and Th2 cell differentiation and Th17 cell differentiation (Fig 5B). Surprisingly, when we compared DEGs between GF+SLE and GF+PBS groups, the difference enrichment results were only concentrated in systemic lupus erythematosus and proteoglycans in cancer (Fig 5C). Overall, these results indicate that fecal microbiota impacted genes expression on both intestines and spleens in recipients mice, and the function of these DEGs were enriched in metabolism pathways and immune response of inflammation, which were closely related to the pathogenesis of SLE.

### 6. Microbiota of human SLE influences the phenotype of metabolism in colonized germ-free mice

The metabolic abilities of the fecal microbiota from recipient mice were tested by using the PM system (Biolog). The compound in well E7 was histidine. Compared Fig 6A and Fig 6B, we noticed that decarboxylation abilities of fecal microbiota from GF+SLE mice (mice had the similar microbiome composition to SLE patients as previously described,) were much lowered than the fecal microbiota from GF+HC mice (Fig 6C). We used HPLC to detected content of histamine in serum, which was the decarboxylation products by gut microbiota, was significantly increased in GF+SLE mice (Fig 6D). The expression of *Hrh4* gene (coding gene of Histamine type 4 receptor) in GF+SLE group spleens was significantly higher than that in GF+HC group (Fig 6E). The phenotype of metabolism indicated that metabolic activities of histidine by microbiota may related to the flare of SLE.

**Figure 6.**
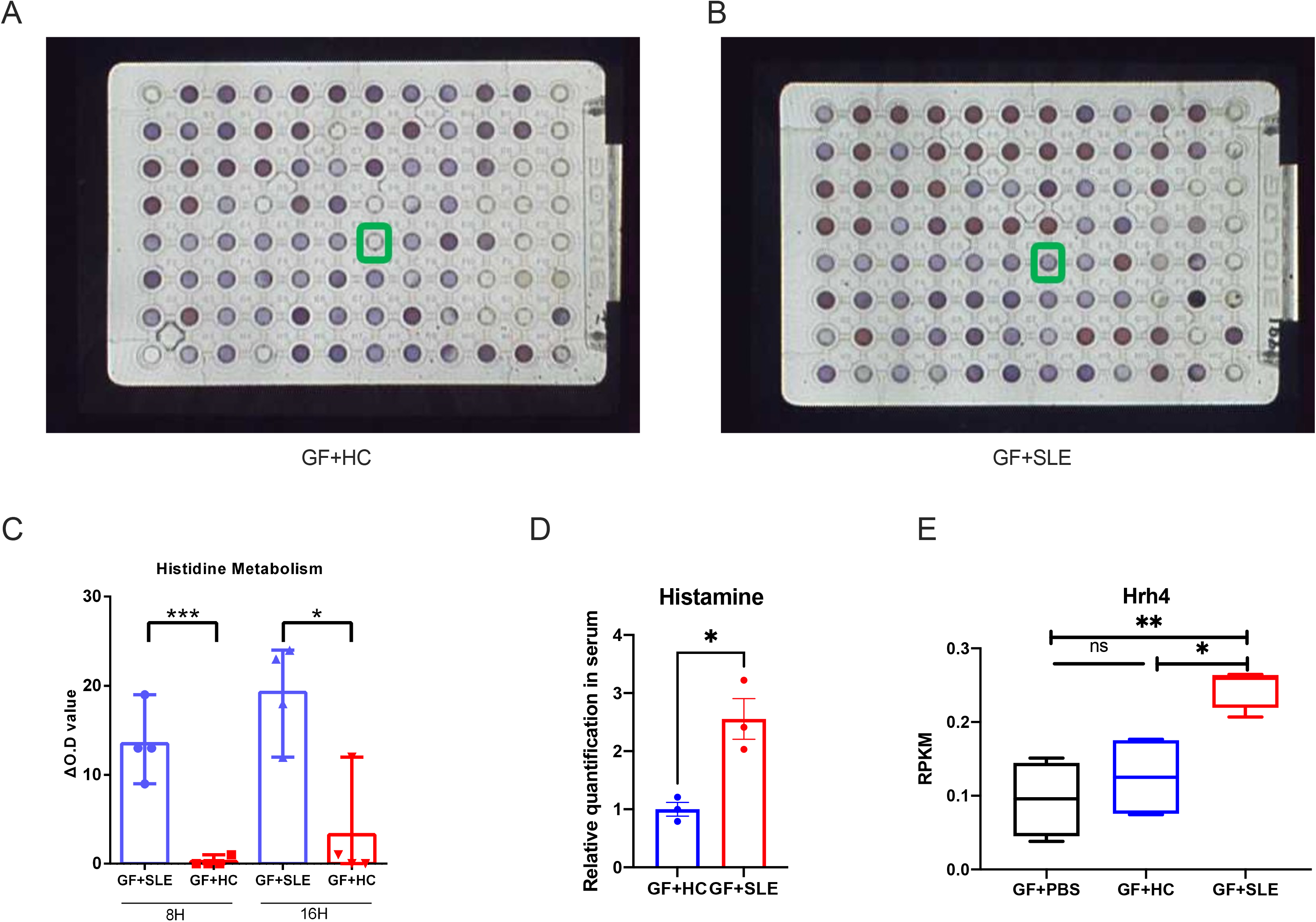
Phenotype of Metabolism in Recipient Mice. (A) Phenotype of metabolic abilities of the fecal microbiota in GF+HC mice. (B) Phenotype of metabolic abilities of the fecal microbiota in GF+SLE mice. (C) Quantification of the metabolic abilities of fecal microbiota from different recipient mice. (D) Relative quantification of histamine in recipient mice serum. (E) RPKM of *Hrh4* gene expression in recipient mice spleen.

## Discussion

Previous studies using integrative biological tools revealed the microbiome alterations present in SLE patients. However, it is not sufficient to identify the causative role of microbiota played without genetic predisposition. Here, we recruited 18 treatment naïve SLE patients along with 7 matched healthy controls, and transferred their fecal microbiota to germ-free C57/B6 mice. The recipients mice developed a series of lupus-like phenotypes, and altered histidine metabolism of the microbiota was observed in SLE recipients mice. In our SLE cohorts, we observed no tangible evidence for reduced gut microbiota diversity in SLE patients, but did observe a differential distribution of bacterial taxa, with an abundance of *Turicibacter* genus, and some *Lactobacillus* species. These bacterial taxa exhibited an expansion in other research of SLE patients [13]. In LEfSe analysis, we noticed that there were no biomarker microbiota for SLE patient group from the kingdom to the genus level, while the only one biomarker of SLE at a species level was a non-dominant species. These results suggest that specific microorganisms play an important role in maintaining healthy homeostasis. However, specific species are not significant biomarker in SLE. A possible explanation for this might be that composition of microbiota is not a key determinant of SLE, while the function of the flora is crucial.

Furthermore, we transplanted aliquots of individual fecal samples into germ-free mice and characterized the representation of the microbiota colonizing residents, SourceTracker results showed that microbiota from human donors contributed more than 75% to communities in their recipient mice, as the communities in recipients had great similarity to their donors. Importantly, the significantly different species maintained the same differences and trends as found in the individual donors. Some microbial species in *Clostridia* order and *Lachnnospiraceae* family were biomarker species (LDA score>4) for GF+HC mice, as these species were identified as a characteristic biomarker for HC group and were significantly different in GF+HC mice as compared with GF+SLE mice.

These results provide us with clues that some special microbial species may be important for healthy gut microbiota. Notably, *Lactobacillus reuteri* was significantly enriched in both SLE patients and recipient mice in our study. *L. reuteri* has been reported to become translocated from gut to the peripheral organs resulting in systemic immune activation; and it can drive autoimmunity in a Toll-like receptor 7-dependent mouse model of SLE [13]. *Ruminococcus gnavus (*of the family *Lachnnospiraceae)*, which represents a symbiont within the healthy human intestinal microbiota, was significantly different between the groups of recipient mice. *R*. *gnavus* species showed a modest enrichment in mice that received SLE fecal microbiota, and this species has been reported to be increased in several diseases including SLE, as expansions of *R. gnavus* in SLE patients has been positively associated with high disease activity, especially in patients with lupus nephritis [14]. In addition, *Turicibacter* was identified as biomarker genus in GF+SLE group (LDA score=4.8), which was also the significantly different species between SLE patients and HC group. Together these results suggested that some special species may play pathogenetic roles in the pathogenesis of SLE.

Two weeks after FMT, we detected significantly increased serum anti-dsDNA antibodies in GF+SLE mice; the increased anti-dsDNA antibodies continued when the recipients were sacrificed at three weeks after FMT. GF mice received SLE donor feces had significantly increased serum antinuclear antibodies (ANA) and total IgG antibodies. Additionally, a series of proinflammatory cytokines including IL-6, IL-8, TNF-α, IL-1β and IFN-γ were significantly increased, while IL-10 was decreased. IL-10 is the prototypical anti-inflammatory cytokine, most classically associated with Foxp3^+^ Tregs [15]. The increased IFN-γ and IFN-α suggested an activation of IFN-type I signature, which is a classic pathway of the pathogenesis of SLE [16, 17]. These phenotype are seen during lupus disease progression, and flares of proinflammation and immune dysregulation. We also noticed that in the colonized mice these patterns were relatively mild compared most models of spontaneous murine lupus. According to these data, we can infer that microbiota trigger lupus associated pathways, which may be reflected as an imbalance of immune inflammation. We also observed a decreased IgA level in SLE fecal samples, although higher fecal IgA levels have been reported in established SLE [14]. After FMT, there were significant increases in IgA in both the recipients with SLE or HC fecal microbiota, while the IgA level were similar between the recipient mice with SLE or HC fecal microbiota. IgA is essential for maintaining gut homeostasis through protection of the mucosal surface from pathogens using mechanisms such as direct binding to commensal bacteria, which together with the mucous layer prevent bacteria from breaching the intestinal epithelial barrier [18]. Moreover, some pathogens or opportunistic commensals may elicit strong intestinal IgA responses in the context of inflammation or dysbiosis [19]. We should also consider that in our model system secretory IgA production was likely independent of T cells. From the content quantity results of IgA antibodies, we noticed that the quantity of IgA antibody in recipients is much lower than that in SLE patients and healthy control, which suggested a milder lupus-like effect was transferred from the donor to these otherwise healthy (i.e., non autoimmune prone) recipient mice. These results indicated that the role of microbiota without genetic susceptibility of SLE was limited.

Lymphocyte distributions in the LP and SP of recipient germ-free mice were different, as shown in Fig 4. B cells and plasma cells in GF+SLE group were significantly increased in both the LP and SP. The increase of Th17 cells and decrease of Tregs were also observed in SP of the mice transplanted with SLE fecal microbiota. Other proinflammation immunocytes including macrophage, B1 cells, immature B cells, geminal center B cells and CXCR3^+^ CD4^+^ T cells, which are involved in both innate and adaptive immunity; CXCR3 ligands are known to play important role on T cells and are key drivers of immune function in the context of autoimmunity [20]. These immune cells were differentially increased in recipient mice with different fecal microbiota. Our results showed that both the adaptive and innate immune systems became altered. The different distribution of immune cells showed that the immune response and microbiome were correlated, which suggests that the changes of lupus-like immune systems(response) in GF+SLE mice can be triggered by the gut microbiome. These findings support the notion that the bacterial community that colonizes the intestine is a major influence on the localized immune response and the peripheral immune response which represent potential mechanism for SLE initiation.

We used transcriptome sequencing to compare changes in intestinal gene expression after FMT. The number of differentially expressed genes (DEGs) showed similar quantity level in different recipients groups, the DEGs number in all recipients groups were more than 25,000. However, the expression of these DEGs in different fecal microbiota recipients mice were greatly different. Comparisons of the function of DEGs in colons between GF+SLE and GF+HC showed the significantly different results were mostly in metabolomic pathways, which indicated that microbiota colonization influenced the gene expression regulating the metabolome of the recipient mice. The DEGs function results in recipients spleens were mainly in immune response pathways. It is somewhat surprising that the most significantly different function of DEGs in GF+SLE mice was systemic lupus erythematosus as it showed in Fig 5. These results strongly suggested fecal microbiota could induce alterations of gene expression in SLE. A possible explanation for these above described results may be that diverse microbiome taxa influenced a range of changes of gene expression; fecal microbiota from SLE patients affect the recipients’ metabolism and immune response, which further contributes to SLE flares.

According to the DEGs results in intestines, we tested the metabolomic phenotype of fecal microbiota from GF+SLE and GF+HC mice. Microbial-associated metabolites serve as a source of immunomodulation [21]. We identified a significant difference in histamine metabolic ability of fecal microbiota from GF+SLE mice. An altered distribution of histidine metabolism has been consistently observed in other studies of SLE patients [22, 23]. Here, we found a more active decarboxylation process of fecal microbiota from GF+SLE mice, which means an increase content of histamine and its metabolites. The biological effect of histamine depends on the type of its receptor. Investigation into the complex regulation function of histamine is still in its infancy. The histamine content in serum is higher in patients with rheumatoid arthritis [24] and elevated in inflamed tissue in atopic dermatitis [25]. In addition, histamine content is significantly increased in the inflamed skin of MRL/lpr mice[26]. Our study showed that GF+SLE mice had significantly increased splenic mRNA expression of the *Hrh4* gene, which encodes for the histamine H4 receptor (H4R). H4R is largely expressed in lymphocytes and it has also been implicated as a regulatory factor in the immune system, which plays roles in inflammatory disorders[27]. Our results support a contribution of intestinal microbiota to histidine metabolism as potential pathway involved in the pathogenesis of SLE.

There are several limitations to this study. Although we have linked gut microbiota to altered histidine metabolism in the pathogenesis of SLE via FMT from SLE patients to GF mice, there is still a lack of direct causal relationship in SLE patients. Further, we have not tested whether altering histidine metabolism, or its product histamine, would affect the flare of SLE. Mechanistically, we have not identified the phenotypic changes in response to variations in histamine or H4R functionally in the recipient mice, we have not yet identified the molecular pathway(s) by which this occurs.

## Conclusion

SLE is associated with alterations of gut microbiota. Fecal microbiota from SLE patients can induce a series of lupus-like phenotypes and altered histidine metabolism in recipient germ-free mice. To mechanistically clarify the role of altered histidine metabolism as a feature of the microbiota trigger for the pathogenesis of SLE, additional experiments will be necessary. In any case, our study identifies a potential molecular pathway target for the treatment of SLE.

## Methods

### 1. Patients information and sample collection

A total of 18 female patients diagnosed as SLE according to the 2010 American College of Rheumatology(ACR) classification criteria and 7 healthy volunteers were recruited for this study from January 2019 to June 2019 at Peking Union Medical College Hospital (Beijing, China) and Renji Hospital (Shanghai, China). The patients’ characteristics, including age, gender, routine clinical blood test results, autoantibody test results, immunoglobin levels, serum complement, and other clinical data were recorded (supplementary table S1). The exclusion criteria were as follows: volunteers with other autoimmune diseases(such as rheumatic arthritis, systemic sclerosis, ankylosing spondylitis and Sjogren’s syndrome et al); patients who received any anti-inflammatory treatment within 1 month before sampling; all participants including healthy and SLE patients who received antibiotics or probiotics treatment in 1 month; all participants with smoking or drinking habits, and all participants who had eaten fermented food, such as kimchi, within 1 week. Clinical data were collected within 3 days before the fecal sample collection. All healthy participants were the same gender with SLE patients (female). They also had comparable age with SLE patients. None of the healthy volunteers had a history of any autoimmune diseases. All samples and clinical information were obtained under the condition of informed consent. This study was conducted with the approval of the Institutional Review Board of Peking Union Medical College Hospital and Renji Hospital in accordance with the Declaration of Helsinki.

The samples, including stool and serum from the same volunteer, were collected at the same day. Stool samples were self-collected after defecation at hospital and immediately placed in the ice. Then the samples were transferred to the laboratory, divided into two parts and put into two frozen tubes. All the stool samples were then transferred to −80°C for long term storage.

### 2. Selection of SLE patient and healthy donors for fecal microbiota transplantation to germ free recipient mice

Stool samples were obtained from 18 patients with SLE and 7 healthy control individuals. The patients were diagnosed according to classification diagnostic criteria revised by the American Society of Rheumatology (ACR) in 2010. All donors were un-treated active SLE patients. None had chronic infectious diseases or current concurrent infections, tumors, pregnancy or lactation. History of surgery and traumatic stress in the past three months were ruled out. Stool samples were *cold chain* transported at anaerobically to the laboratory and frozen at −80℃. Diluted sample was gavaged 5 times into 9-week-old germ free C57/B6J mice (200μL/time per mouse, one mouse per donor) with 2 days interval. Mice in GF+HC group were gavaged with fecal from healthy donors, and mice in GF+SLE group were gavaged with fecal from SLE patients. A blank control group (GF+PBS) was gavaged 5 times with PBS solution (200μL per mouse). After 3 weeks, mouse serum autoimmune antibodies and fecal IgA antibodies were assessed. Mice were sacrificed thereafter, and samples were collected.

### 3. Fecal Microbiota Transplantation

To prepare the stool for gavage, stools from healthy donors were mixed at equal weight, samples from SLE patients were separated individually. One gram of the sample or mixed stool was then suspended in 5 mL of PBS and vortexed thoroughly. An aliquot of 200 μL suspension was used for gavage into each mouse.

Germ free (GF) mice on a C57/B6J background were generated and provided by Institute of Laboratory Animal Sciences (ILAS), a member of the American Association for the Accreditation of Laboratory Animal Care-accredited facilities. Studies were performed in accordance with the guidelines of the Institutional Animal Care and Use Committees of ILAS. Adult GF mice were divided into three groups with each 5 mice. The blank control group, GF+PBS, were gavaged with phosphate-buffered saline (PBS) solution. The control group, GF+HC, were gavaged with fecal from healthy donors. The GF+SLE group were gavaged with fecal from SLE patients. All recipient mice were gaveged 5 times, one day interval and lasted for ten days. All recipient mice were sacrificed at week 4 after transplantation.

### 4. 16s rDNA sequencing for fecal bacterial profiles and analysis

The feces of healthy donors and SLE patients were collected in sterile tube, transported cold chain and quickly put into −80° freezer. Feces of recipients were collected at the 2^nd^ week and 3^rd^ week after transplantation. otal genome DNA from samples was extracted using CTAB/SDS method. DNA concentration and purity was monitored on 1% agarose gels. According to the concentration, DNA was diluted to 1ng/μL using sterile water. PCR amplification of V4 regions of bacterial 16S rRNA gene amplicons for paired end sequencing (2 x 250 bps) on the IonS5^TM^ XL platform was performed using universal primer sequences (515F, 5’-GTGCCAGCMGCCGCGGTAA-3’; 806R, 5’GGACTACHVGGGTWTCTAAT-3’). PCR products was mixed in equidensity ratios. Then, mixture PCR products was purified with GeneJETTM Gel Extraction Kit (Thermo Scientific). Sequencing libraries were generated using Ion Plus Fragment Library Kit 48 rxns (Thermo Scientific) following manufacturer’s recommendations. The library quality was assessed on the Qubit@ 2.0 Fluorometer (Thermo Scientific). Lastly, the libraries were sequenced on an IonS5™XL platform and 400bp/600 bp single-end reads were generated.

Paired-end reads were assigned to samples using Cutadapt based on their unique barcode and truncated by cutting off the barcode and primer sequence; Reads were merged using FLASH (V1.2.7)[28].Quality filtering on the raw reads were performed under specific filtering conditions to obtain the high-quality clean reads according to the Cutadapt quality-controlled process. The reads were compared with the reference database using UCHIME algorithm to detect chimera sequences, and then the chimera sequences were removed[29, 30]. Then, the clean reads were finally obtained. Sequences analysis were performed by Uparse software (Uparse v7.0.1001)[31]. Sequences with ≥97% similarity were assigned to the same OTUs. Representative sequence for each OTU was screened for further annotation. For each representative sequence, the Silva Database was used to annotate taxonomic information. In order to study phylogenetic relationship of different OTUs, and the difference of the dominant species in different samples (groups), multiple sequence alignment were conducted using the MUSCLE software (Version .8.31)[32]. OTU abundance information were normalized using a standard of sequence number corresponding to the sample with the least sequences. Subsequent analysis of alpha diversity and beta diversity were all performed basing on this output normalized data. Alpha diversity and Beta diversity indices in our samples were calculated with QIIME (Version1.7.0) and displayed with R software (Version 2.15.3). Differential abundance analysis was performed using all-against-all detection algorithm of LEfSe program. We filtered differentially abundant OTUs by false discovery rate-adjusted (FDR) significance of ≤ 0.05 from linear discriminant analysis (LDA). Average robust fold change for each OTU was computed using the fcros package in the R Project for Statistical Computing, by pairwise sample comparison with the default quartile feature exclusion criteria based on fold change rank-order statistics. All sequencing and analysis were performed by Novogene, Beijing, China.

### 5. Enzyme-Linked Immuno Sorbent Assay (ELISA) for serum antibodies and cytokines detection

ELISA were performed to measure the protein levels of anti-dsDNA antibodies, anti-ANA antibodies and total IgG antibodies in mouse serum; IL-6, IL-8, IL-10, TNF-α, IFN-α and IFN-γ in mouse serum. ELISA KITs (MEIMIAN, Jiangsu Province, China) were commercially obtained, and detection was performed following the manufacturer’s instructions.

### 6. ELISA for fecal IgA antibodies

Feces from recipients mice were collected at the 3^rd^ week after transplantation. For each recipient mouse, 0.1g feces was suspended in 0.9ml sterile PBS solution (pH 7.4) and vortexed thoroughly. Centrifuge the mixture at 2000rpm, 10min. The supernatant was obtained for detection. The ELISA KIT (MEIMIAN, Jiangsu Province, China) was commercially available, and all detection were performed following the manufacturer’s instructions.

### 7. Isolation of splenic lymphocytes and Lamina Propria lymphocytes (LPL) for flow cytometry

Spleens were removed and cut into pieces in cold PBS. After filtration through a 200-gauge steel mesh and the removal of red blood cells with RBC lysis buffer (BD Biosciences, USA), splenocytes were collected for further experimentation. The isolation of small intestine LPLs was performed as previously described [33].

### 8. Flow cytometry

Splenocytes and LPLs were incubated with a rat anti-mouse CD16/CD32 mAb (BioLegend, USA) for 30 min to block Fc receptors. Then, the cells were stained on ice with fluorochrome-conjugated monoclonal antibodies for 30 min as previously described [34]. To perform intracellular staining, cells were treated using a Fixation/Permeabilization Solution kit (BD Pharmingen™, USA) and then stained with the appropriate antibodies. The antibodies used in this study are listed in supplementary file table S2. Cells were assayed with a Symphony A5 cytometer (BD Biosciences), and the resulting data was analysed using FlowJo (ver. 10.4).

### 9. RNA extraction and transcriptome sequencing

The colon and spleen tissues of GF mice were collected and stored at − 80°immediately after euthanization. Total RNA was isolated using the TRIzol reagent (Invitrogen, CA, USA), and 3 μg RNA per sample was used for analysis. Sequencing libraries were generated using NEBNext® UltraTM RNA Library Prep Kit for Illumina® (New England Biolabs, USA) following manufacturer’s recommendations, and index codes were added for attribution of each sample. The library was sequenced on an Illumina Hiseq platform, and 125/150 bp paired-end reads were generated. Raw data (raw reads) in the fastq format were first processed through in-house Perl scripts. Clean data (clean reads) were then obtained by removing reads containing adapter sequences, poly-N sequences, and low quality reads from the raw data. Reference sequences were from STAR, and paired-end clean reads were aligned to the reference genome using STAR (ver. 2.5.1b). HTSeq (ver. 0.6.0) was used to count the read numbers for each gene. The number of fragments per kilobase of transcript per million mapped reads of each gene was calculated based on the length of the gene and the number of reads mapped to that gene. Differential expression genes in the two groups were determined by using the DESeq2 in R software. The Cluster Profiler in R software was used to test the statistical significance of enrichment of different genes, based on pathways in the Kyoto Encyclopedia of Genes and Genomes. Gene Ontology (GO) enrichment analysis of differentially expressed genes was implemented using the cluster Profiler in R software, with correction for gene length bias. The differential expression of genes was analyzed by multiple hypothesis testing, based on the principle of negative binomial distribution. A significant difference is presented as a false-discovery rate (FDR) adjusted p-value, is also called a q-value. All sequencing and analysis was performed by Novogene, Beijing, China.

### 10. Phenotype of metabolism (PM) analysis and metabolic profile

Stools from recipient mice were collected at the 3^rd^ week after transplantation. For each recipient mouse, 0.1g feces was suspended in 0.9ml sterile PBS solution (pH 7.4) and vortexed thoroughly. Centrifuge the mixture at 1200rpm, 10min. The supernatant was obtained for detection. Metabolomic profiling of the fecal microbiota were performed using Phenotype MicroArray (GEN III) microplates. For each well, 200ul supernatant were loaded and followed by incubation in Biolog’s Omnilog PM instrument at 37°C. The OmniLog reader was set to measure the tetrazolium reduction, at 15-minute intervals for 24 hours. The data collected by the OmniLog® Phenotype MicroArray (PM) system was organised using the OmniLog® PM software (Biolog, USA). Based on the manufacturer’s instructions. Values above 0.2 were considered positive, meaning a given bacteria were able to use a given substrate, and values below 0.2 were considered negative[35]. Data analysis for Biolog PM results was executed in Kinetic Analysis program and Graphpad Prism.

### 11. UPLC–MS/MS analysis of serum

For serum samples, 400 μ L of 80% methanol water solution (4 times the volume of methanol) was added to 100μL of serum sample in a EP tube, vortex oscillation, standing at - 20 ° C for 60 min, 14000 g, centrifugation at 4 ° C for 20 min, taking a certain amount of supernatant into 1.5 ml centrifuge tube, vacuum freeze-drying, the residue was dissolved in 100μL complex solvent, vortex oscillation, and centrifugation at 14000 g and 4 ° C for 15 min. the supernatant was injected for LC-MS analysis. An ultra-performance liquid chromatography coupled to tandem mass spectrometry system was performed by Novogene, Beijing, China. The optimized instrument settings and analytical quality control procedures are briefly described in supplementary file.

### 12. Statistics analysis

Mean and SEM were calculated by averaging the results of three independent experiments performed with samples obtained from individual mouse for each experiment. Prism software (GraphPad, San Diego, CA) was used for statistical analysis with the unpaired t-test for comparison between two groups and the one-way ANOVA with *Tukey* multiple comparison for multiple groups. P values p < 0.05 were considered significant. ∗ represent p < 0.05. ∗∗ represent p < 0.01. ∗∗∗ represent p < 0.001.

## Declarations

### Ethics approval and consent to participate

The study was approved by local ethics committees (Peking Union Medical College Hospital, Renji Hospital), and informed consent was obtained from all subjects.

All animal protocols were approved by Ethics Committee for Animal Care and Use at the Institute Laboratory Animal Science, Chinese Academy of Medical Sciences (IACUC No. NHT17002).

### Consent for publication

Not applicable.

### Availability of data and material

The datasets used in the current study are available from the corresponding author on reasonable request.

### Authors’ contributions

Substantial contributions to the conception or design of the study or the acquisition, analysis, or interpretation of data used in the study: YM, RG, ML, LL and HN. Participants’ faeces collection: RG, YS, ZL. Drafting the study or revising it critically for important intellectual content: YM, GJS and HN. Assistance with the design of the study: QW, JY, LH, GC and XL. Final approval of the version to be published: YM, ML, LL and HN. All authors read and approved the final manuscript.

## Acknowledgements

The authors thank Dr. Shuangyue Zhang, Guixiang Zou and Xiaoxue Xu for technical support. The authors thank Dr. Xi Yang and Dr. Rex Wei Wang for excellent technical support in flow cytometry panel design.

## Abbreviations

ACR: American College of Rheumatology
Anti-dsDNA: Anti-double stranded DNA
ANA: anti-nuclear antibodies
DEGs: Differential expression of genes
ELISA: Enzyme-linked immunosorbent assay
FMT: Fecal microbiota transplantation
GC: Germinal center
GF: Germ free
GO: Gene ontology
H4R: histamine H4 receptor
ILAS: Institute of Laboratory Animal Sciences
IBD: Inflammatory bowel disease
LDA: linear discriminant analysis
LEfSe: LDA Effect Size
IRFs: IFN regulatory factors
LN: Lupus nephritis
LP: Lamina propria
MRL/lpr: MRL/Mp-Fas lpr mouse
NGS: Next-generation sequencing
OTUs: Operational taxonomic unit
PBS: Phosphate-buffered saline solution
PBs: Plasma blast cells
PCA: Principal component analysis
PCs: Plasma cells
qRT-PCR: quantitative real-time PCR
RA: Rheumatoid arthritis
SLE: Systemic lupus erythematosus
Tnf: tumor necrosis factor
Treg: Regulatory T cells

---

**Figure 1S.**
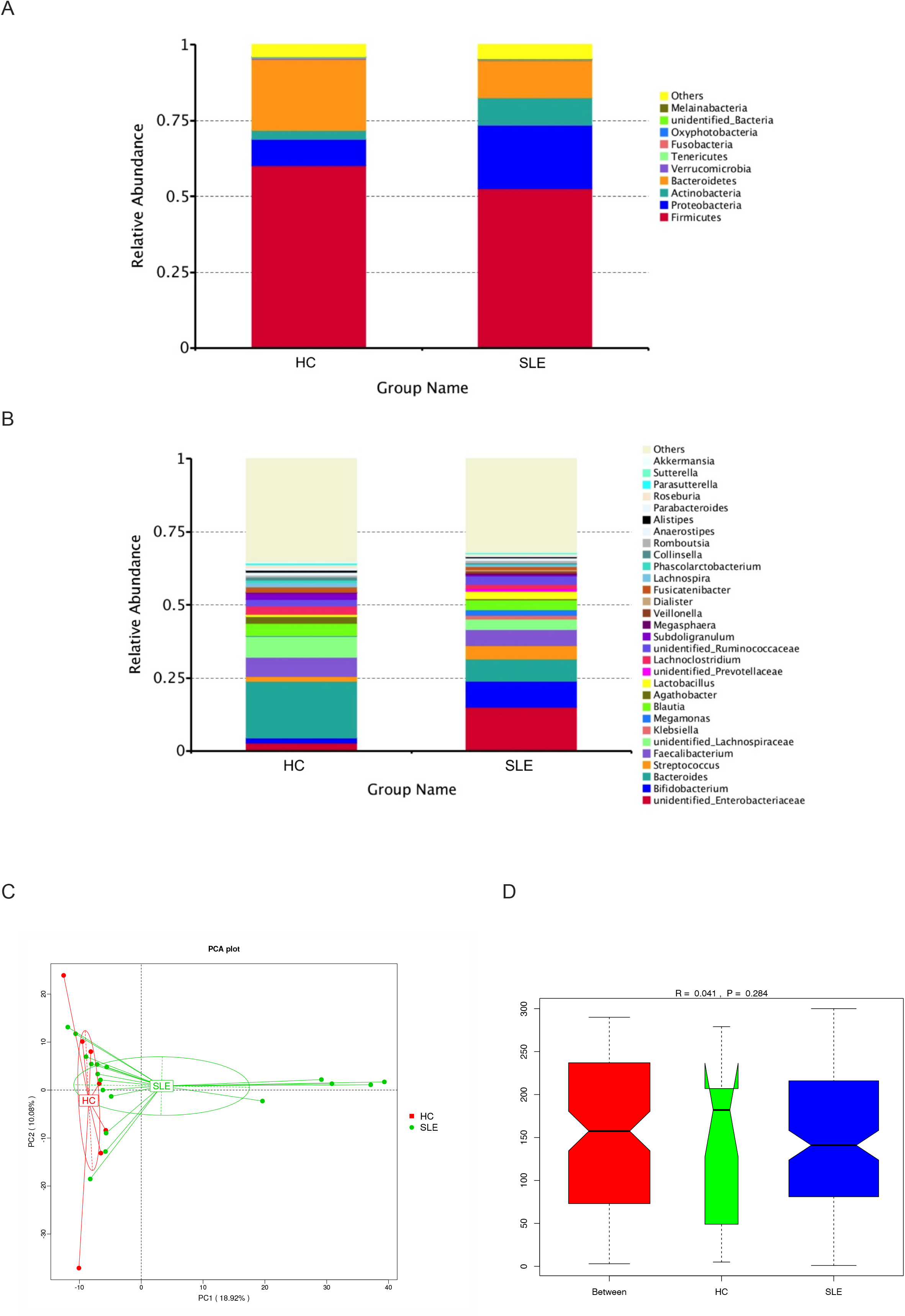
Analysis of gut microbiota in SLE patients and healthy control. Top 10 bacterial phylum(A) and Top 30 bacterial genus(B) of SLE patients and healthy control. PCA analysis of two groups (C). Anosim analysis result between two groups (D).

**Figure 2S.**
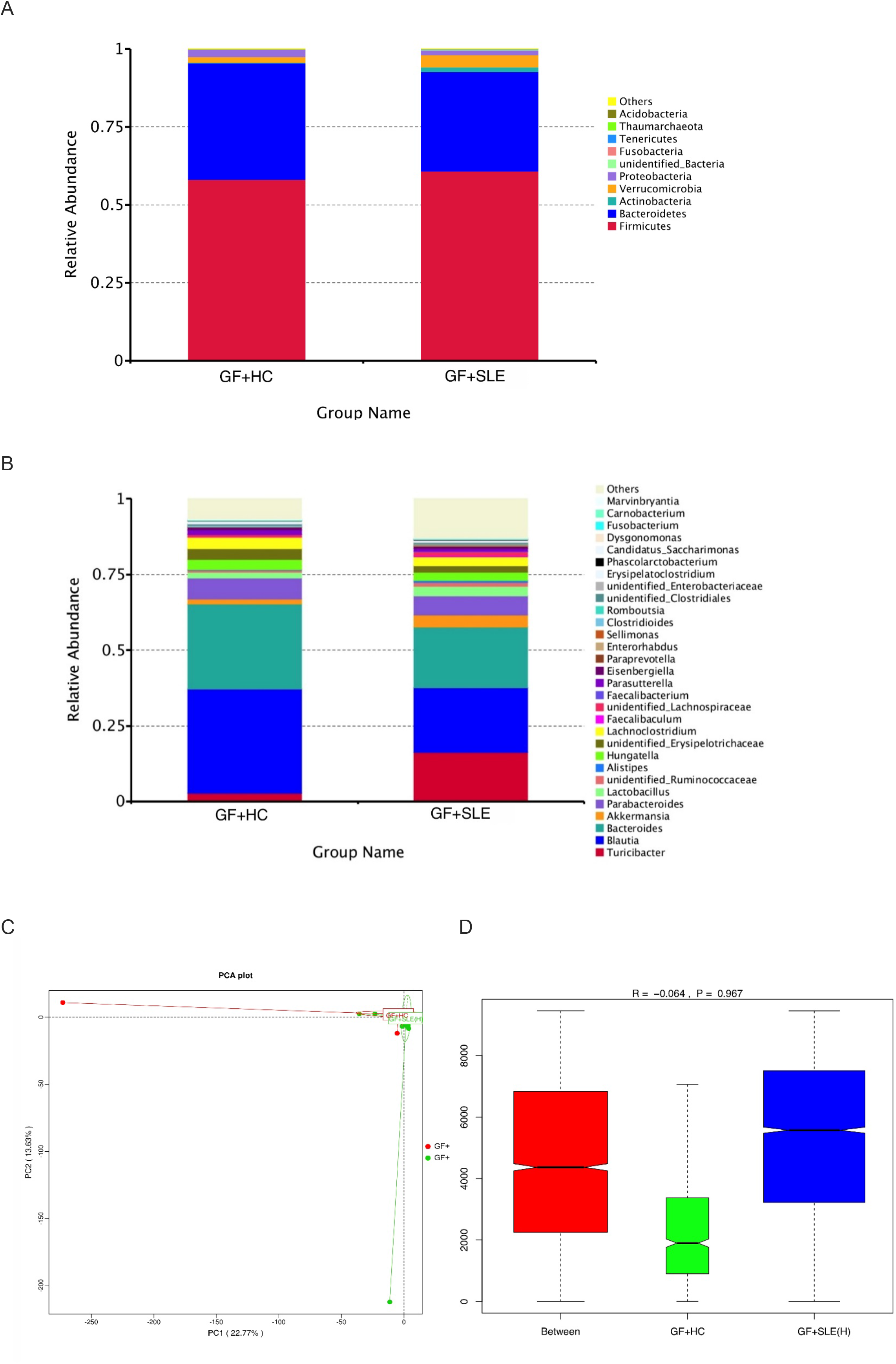
Analysis of gut microbiota in GF+SLE group and GF+HC group. Top 10 bacterial phyla (A) and Top 30 bacterial genera (B) of GF+SLE group and GF+HC group. PCA analysis of two recipient groups (C). Anosim analysis result between the two recipient groups (D).

**Figure 4S.**
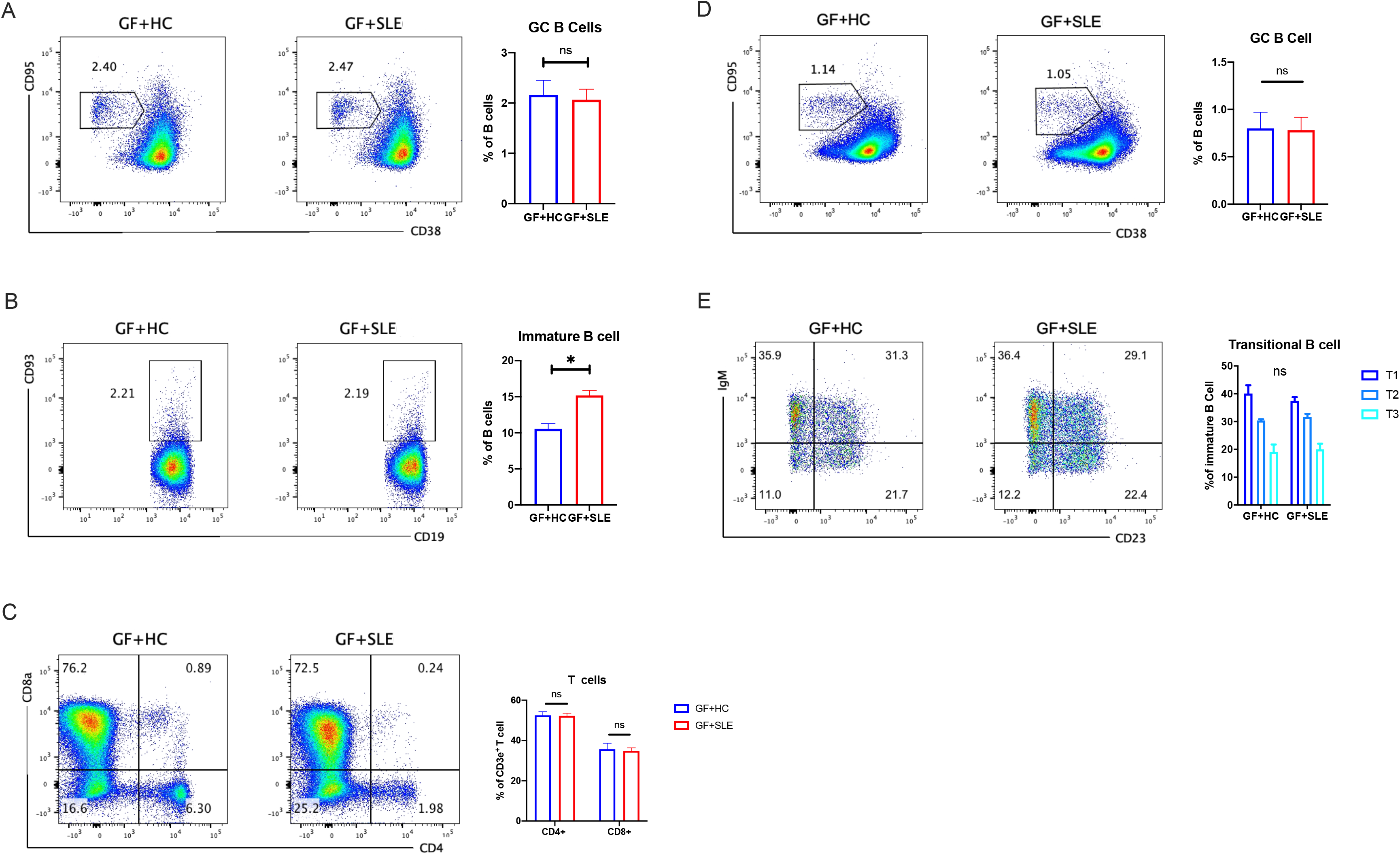
Flow cytometry analysis of the distribution of lymphocytes in the lamina propria (LP) and spleen (SP) of GF+HC mice and GF+SLE mice. (A) Plots of germinal center B cells (GC B cells, left) and percentages of GC B cells in LP (4 mice per group, right). (B) Plots of immature B cells (left) and percentages of immature B cells in LP (4 mice per group, right). (C) Plots of CD4+ and CD8+ T cells (left) and percentages of CD4+ and CD8+ T cells in LP (4 mice per group, right). (D) Plots of germinal center B cells (GC B cells, left) and percentages of GC B cells in SP (4 mice per group, right). (E) Plots of transitional B cells (left) and percentages of transitional B cells in SP (4 mice per group, right). Data represent one of two independent experiments. A Mann-Whitney t-test was used to compare groups, and error bars represent means ± SEMs. *P < 0.05.

**Figure 5S.**
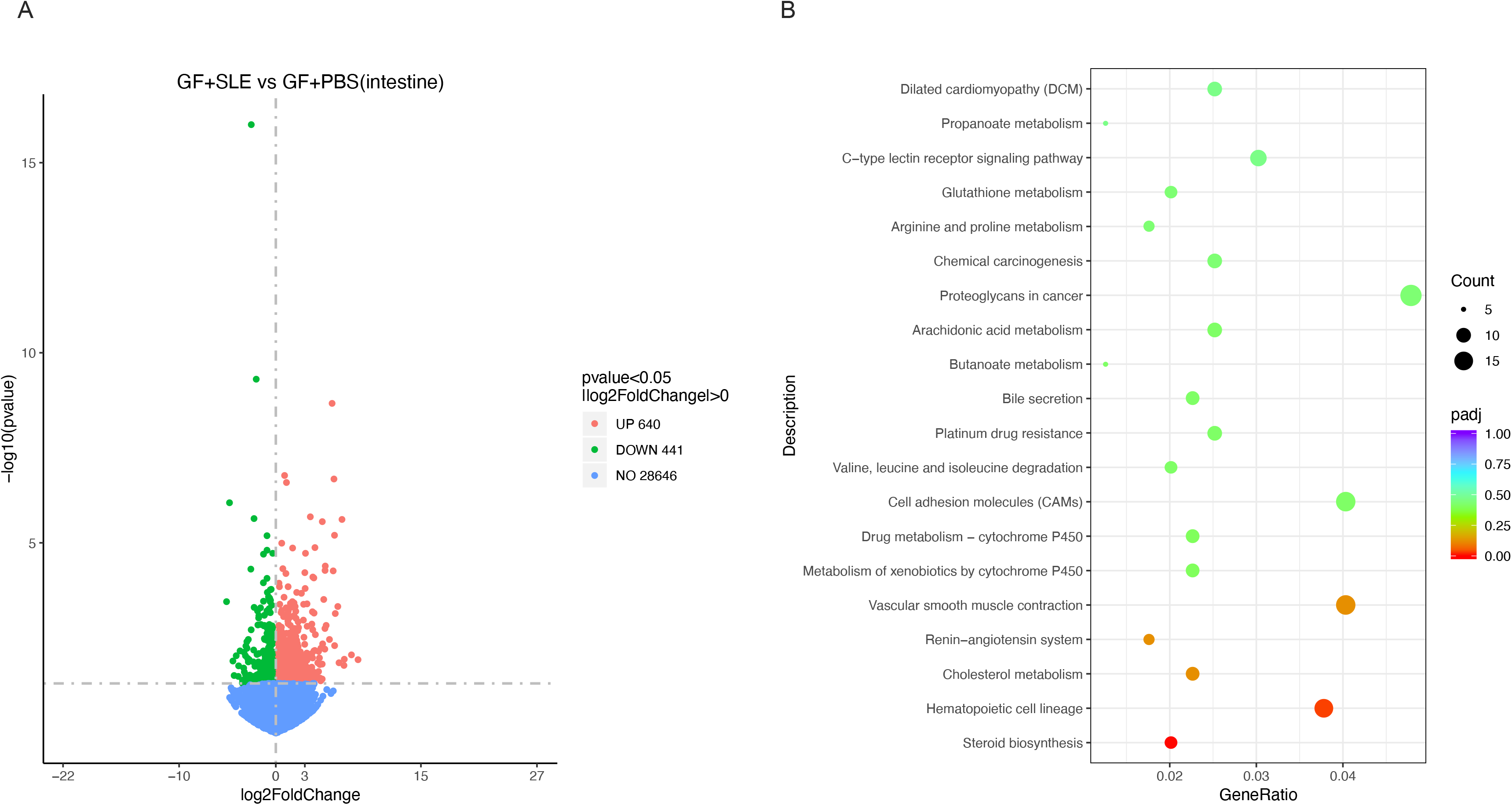
Differential expression of genes (DEGs) in the intestines of GF mice after different treatments. Comparison of the GF + SLE and the GF + PBS groups. The numbers of differential expressed genes in colons showed in volcano plot (A) and its enrichment in KEGG (B).

UPLC-MS/MS instrument settings

UPLC-MS/MS instrument settings
| **UPLC** |  |
| --- | --- |
| Column Temp. (°C) | 40 |
| Sample Manager Temp. (°C) | 10 |
| Mobile Phases | Positive mode A= formic acid (0.1%), acetonitrile (95%) and ammonium acetate(10mM); and B= formic acid (0.1%),acetonitrile (50%) and ammonium acetate(10mM); Negative mode A= acetonitrile (95%), ammonium acetate(10mM), pH=9; and B= acetonitrile (50%), ammonium acetate(10mM), pH=9. |
| Gradient Conditions | 0-1 min (2% B), 1-17 min (50% B), 17-17.5 min (50% B), 17.5-18 min (2%B), 18-20 min (2%B) |
| Flow Rate (mL/min) | 0.30 |
| Injection Vol. (μL) | 5.0 |
| **MASS SPECTROMETER** |  |
| Spray Voltage (Kv) | 3.2 (ESI-) |
| Capillary Temp (°C) | 320 |
| Sheath gas flow rate | 35 arb |
| Aux Gas flow rate | 10 arb |

The offline data (. raw) file is imported into the CD search software, and the retention time, mass charge ratio and other parameters are calculated. The peaks of different samples were aligned according to the retention time deviation of 0.2 min and mass deviation of 5 ppm to make the identification more accurate. The peaks were extracted according to the set mass deviation of 5 ppm, signal intensity deviation of 30%, signal-to-noise ratio of 3, minimum signal intensity of 100000, additive ions and other information, the peak area was quantified, and the target ions were integrated. The molecular formula was predicted and compared with mzCloud database. The background ions were removed with blank samples, and the quantitative results were normalized with QC samples. Finally, the data identification and quantitative results were obtained.

**Supplementary table S1.** NO.1-4 are FMT donors

| No. stool | Gender | Age | BMI | SLEDAI | Manifestation | ANA |
| --- | --- | --- | --- | --- | --- | --- |
| 1* | F | 25 | 19.83 | 4 | malar rush,abdominal pain, diarrhea, constipation | (+)H1:1280 |
| 2* | F | 31 | 21.83 | 21 | abdominal pain,micturition and urgent emiction | (+) S1:1280 |
| 3* | F | 24 | 24.75 | 14 | Abdominal pain, proteinurine, malar rush, erythra | S1:640 |
| 4* | F | 33 | 21.91 | 8 | Edema | H1:1280 |
| 5 | F | 32 | 21.67 | 11 | Arthralgia, fever, hemorrhinia | H1:640 |
| 6 | F | 52 | 25.07 | 10 | Abdominal pain, diarrhea, abdominal distention | S1:80 |
| 7 | F | 21 | 21.09 | 21 | Edema, fever | H1:320 |
| 8 | F | 28 | 21.72 | 8 | Petechia, unconsciousness, fever | HS1:160 |
| 9 | F | 46 | 26.37 | 10 | Edema, anxious, erythra, urine abnormal | H1:80 |
| 10 | F | 18 | 27.93 | 12 | malar rush, arthralgia, edema, skin blisters | (+)S1:640 |
| 11 | F | 30 | 26.61 | 18 | fever,edema,figertip erythra, | (+) H1:320 |
| 12 | F | 13 | 17.48 | 18 | fever,rash,chest pain,diarrhea, | (+) H1:1280 |
| 13 | F | 28 | 26.23 | 12 | fever,arthralgia,edema. | (+) H1:640 |
| 14 | F | 57 | 20.28 | 8 | rash,edema, foam urine | (+) S1:80 |
| 15 | F | 25 | 22.21 | 10 | fever,rash,arthralgia,edema, foam urine | (+) H1:1280 |
| 16 | F | 41 | 19.53 | 9 | rash,arthralgia,fever,headache | (+) SN1:320 |
| 17 | F | 44 | 24.24 | 10 | vision loss, edema, abdominal pain, diarrhea, oral ulcer | (+) S1:160 |
| 18 | F | 23 | 16.16 | 10 | Visual field defected and loss, headache,edema,diarrhea, limb | (+) H1:320 |

**Supplementary table S2.**

| anti-dsDNA ELISA IU/ml | C3 g/l | C4 g/l | IgG g/l | IgA g/l | IgM g/l | Infection | Abx | Surgery | Tumor | DM | other immune disease |
| --- | --- | --- | --- | --- | --- | --- | --- | --- | --- | --- | --- |
| 279 | 0.360 | 0.005 | 7.16 | 1.37 | 1.31 | N | N | N | N | N | N |
| 162 | 0.399 | 0.134 | 29.67 | 5.83 | 1.36 | N | N | N | N | N | N |
| 248 | 0.268 | 0.021 | 25.46 | 3.08 | 0.8 | N | N | N | N | N | N |
| >800 | 0.741 | 0.109 | 9.84 | 2.84 | 0.98 | N | N | N | N | N | N |
| 196 | 0.298 | 0.082 | 13.34 | 1.24 | 1.79 | N | N | N | N | N | N |
| / | 0.598 | 0.050 | 6.08 | 1.25 | 0.48 | N | N | N | N | N | N |
| 416 | 0.189 | 0.001 | 14.01 | 4.7 | 0.95 | N | N | N | N | N | N |
| / | 0.765 | 0.162 | 10.57 | 1.43 | 0.97 | N | N | N | N | N | N |
| / | 0.832 | 0.133 | 6.58 | 2.38 | 0.19 | N | N | N | N | N | N |
| / | 0.937 | 0.189 | 4.04 | 1.87 | 1.43 | N | N | N | N | N | N |
| 398 | 0.322 | 0.055 | 9.42 | 3.41 | 0.99 | N | N | N | N | N | N |
| >800 | 0.589 | 0.122 | 7.50 | 3.22 | 0.90 | N | N | N | N | N | N |
| 102 | 0.302 | 0.034 | 11.46 | 2.37 | 0.61 | N | N | N | N | N | N |
| — | 0.954 | 0.26 | 7.92 | 2.03 | 0.31 | N | N | N | N | N | N |
| >800 | 0.421 | 0.129 | 6.51 | 2.76 | 1.44 | N | N | N | N | N | N |
| 275 | 0.376 | 0.045 | 11.7 | 1.78 | 0.57 | N | N | N | N | N | N |
| — | 0.388 | 0.088 | 5.26 | 1.7 | 1.62 | N | N | N | N | N | N |
| 406 | 0.630 | 0.152 | 10.14 | 0.68 | 0.45 | N | N | N | N | N | N |

| Marker | Fluorochrome |
| --- | --- |
| CD19 | BUV737 |
| B220 (CD45) | BUV395 |
| CD138 | BV421 |
| CD38 | PE |
| CD95 | BV510 |
| IgM | APC |
| IgD | BV786 |
| PNA | FITC |
| CD93 | BV650 |
| CD3e | APC-Cy7 |
| CD4 | BUV395 |
| CD8a | Percp-Cy5.5 |
| CD25 | BB515 |
| Foxp3 | BV421 |
| RORyt | BV786 |
| Live/Death | FVS700 |

## References

1. Yu, J., et al., Metagenomic analysis of faecal microbiome as a tool towards targeted non-invasive biomarkers for colorectal cancer. Gut, 2017. 66(1): p. 70–78.

2. Liu, H., et al., Alterations in the gut microbiome and metabolism with coronary artery disease severity. Microbiome, 2019. 7(1): p. 68.

3. De Luca, F. and Y. Shoenfeld, The microbiome in autoimmune diseases. Clin Exp Immunol, 2019. 195(1): p. 74–85.

4. Justiz Vaillant, A.A., et al., Systemic Lupus Erythematosus, in StatPearls. 2021: Treasure Island (FL).

5. Hevia, A., et al., Intestinal dysbiosis associated with systemic lupus erythematosus. mBio, 2014. 5(5): p. e01548–14.

6. He, Z., et al., Alterations of the gut microbiome in Chinese patients with systemic lupus erythematosus. Gut Pathog, 2016. 8: p. 64.

7. Chen, B.D., et al., An Autoimmunogenic and Proinflammatory Profile Defined by the Gut Microbiota of Patients With Untreated Systemic Lupus Erythematosus. Arthritis Rheumatol, 2021. 73(2): p. 232–243.

8. Mu, Q., et al., Control of lupus nephritis by changes of gut microbiota. Microbiome, 2017. 5(1): p. 73.

9. Luo, X.M., et al., Gut Microbiota in Human Systemic Lupus Erythematosus and a Mouse Model of Lupus. Appl Environ Microbiol, 2018. 84(4).

10. Mu, Q., et al., Antibiotics ameliorate lupus-like symptoms in mice. Sci Rep, 2017. 7(1): p. 13675.

11. de la Visitacion, N., et al., Gut microbiota contributes to the development of hypertension in a genetic mouse model of systemic lupus erythematosus. Br J Pharmacol, 2021.

12. Ma, Y., et al., Applications of Next-generation Sequencing in Systemic Autoimmune Diseases. Genomics Proteomics Bioinformatics, 2015. 13(4): p. 242–9.

13. Zegarra-Ruiz, D.F., et al., A Diet-Sensitive Commensal Lactobacillus Strain Mediates TLR7-Dependent Systemic Autoimmunity. Cell Host Microbe, 2019. 25(1): p. 113–127 e6.

14. Azzouz, D., et al., Lupus nephritis is linked to disease-activity associated expansions and immunity to a gut commensal. Ann Rheum Dis, 2019. 78(7): p. 947–956.

15. Chen, J. and X.S. Liu, Development and function of IL-10 IFN-gamma-secreting CD4(+) T cells. J Leukoc Biol, 2009. 86(6): p. 1305–10.

16. Elkon, K.B. and A. Wiedeman, Type I IFN system in the development and manifestations of SLE. Curr Opin Rheumatol, 2012. 24(5): p. 499–505.

17. Liu, W., et al., IFN-gamma Mediates the Development of Systemic Lupus Erythematosus. Biomed Res Int, 2020. 2020: p. 7176515.

18. Pabst, O., New concepts in the generation and functions of IgA. Nat Rev Immunol, 2012. 12(12): p. 821–32.

19. Kau, A.L., et al., Functional characterization of IgA-targeted bacterial taxa from undernourished Malawian children that produce diet-dependent enteropathy. Sci Transl Med, 2015. 7(276): p. 276ra24.

20. Karin, N., CXCR3 Ligands in Cancer and Autoimmunity, Chemoattraction of Effector T Cells, and Beyond. Front Immunol, 2020. 11: p. 976.

21. Ost, K.S. and J.L. Round, Communication Between the Microbiota and Mammalian Immunity. Annu Rev Microbiol, 2018. 72: p. 399–422.

22. Ouyang, X., et al., (1)H NMR-based metabolomic study of metabolic profiling for systemic lupus erythematosus. Lupus, 2011. 20(13): p. 1411–20.

23. Yan, B., et al., Serum metabolomic profiling in patients with systemic lupus erythematosus by GC/MS. Mod Rheumatol, 2016. 26(6): p. 914–922.

24. Adlesic, M., et al., Histamine in rheumatoid arthritis. Scand J Immunol, 2007. 65(6): p. 530–7.

25. Schaper-Gerhardt, K., et al., The role of the histamine H4 receptor in atopic dermatitis and psoriasis. Br J Pharmacol, 2020. 177(3): p. 490–502.

26. Tachibana, T., et al., Histamine metabolism in skin of MRL/l mice. Arch Dermatol Res, 1985. 278(1): p. 57–60.

27. Zampeli, E. and E. Tiligada, The role of histamine H4 receptor in immune and inflammatory disorders. Br J Pharmacol, 2009. 157(1): p. 24–33.

28. Magoc, T. and S.L. Salzberg, FLASH: fast length adjustment of short reads to improve genome assemblies. Bioinformatics, 2011. 27(21): p. 2957–63.

29. Edgar, R.C., et al., UCHIME improves sensitivity and speed of chimera detection. Bioinformatics, 2011. 27(16): p. 2194–200.

30. Haas, B.J., et al., Chimeric 16S rRNA sequence formation and detection in Sanger and 454-pyrosequenced PCR amplicons. Genome Res, 2011. 21(3): p. 494–504.

31. Edgar, R.C., UPARSE: highly accurate OTU sequences from microbial amplicon reads. Nat Methods, 2013. 10(10): p. 996–8.

32. Edgar, R.C., MUSCLE: multiple sequence alignment with high accuracy and high throughput. Nucleic Acids Res, 2004. 32(5): p. 1792–7.

33. Weigmann, B., et al., Isolation and subsequent analysis of murine lamina propria mononuclear cells from colonic tissue. Nat Protoc, 2007. 2(10): p. 2307–11.

34. Ma, Y., et al., Gut microbiota promote the inflammatory response in the pathogenesis of systemic lupus erythematosus. Mol Med, 2019. 25(1): p. 35.

35. Lugtenberg, B.J., L.V. Kravchenko, and M. Simons, Tomato seed and root exudate sugars: composition, utilization by Pseudomonas biocontrol strains and role in rhizosphere colonization. Environ Microbiol, 1999. 1(5): p. 439–46.

